# Hemoglobin drives inflammation and initiates antigen spread and nephritis in lupus

**DOI:** 10.1101/2020.11.27.399501

**Authors:** Hritika Sharma, Anjali Bose, Uma Kumar, Rahul Pal

**Author notes:** Corresponding author: Rahul Pal, Immunoendocrinology lab, National Institute of Immunology, Aruna Asaf Ali Marg, JNU Complex, New Delhi – 110067, India.

## Abstract

Hemoglobin (Hb) has well-documented inflammatory effects and is normally efficiently scavenged; clearance mechanisms can be overwhelmed during conditions of erythrocyte lysis, a condition that may occur in systemic lupus erythematosus. Whether Hb is preferentially inflammatory in lupus and additionally induces autoreactivity against prominent autoantigens was assessed. Peripheral blood mononuclear cells derived from SLE patients secreted higher levels of lupus-associated inflammatory cytokines when incubated with Hb, effects negated by haptoglobin. Hb (more particularly, ferric Hb) triggered the preferential release of lupus-associated cytokines from splenocytes, B cells, CD4 T cells, CD8 T cells and plasmacytoid dendritic cells isolated from aging NZM2410 mice, and also had mitogenic effects on B cells. Ferric Hb activated multiple signaling pathways which were differentially responsible for the generation of specific cytokines; inflammatory signaling also appeared to be cell-context dependent. Pull-downs, followed by mass spectrometry, revealed interactions of Hb with several lupus-associated autoantigens; co-incubation of ferric Hb with apoptotic blebs (structures which contain packaged autoantigens, believed to trigger lupus autoreactivity) revealed synergies (in terms of cytokine release and autoantibody production *in vitro*) that were also restricted to the lupus genotype. Infusion of ferric Hb into NZM2410 mice led to enhanced release of lupus-associated cytokines, the generation of a spectrum of autoantibodies, and enhanced-onset glomerulosclerosis. Given that the biased recognition of ferric Hb in a lupus milieu, in concert with lupus-associated autoantigens, elicits the generation of inflammatory cytokines from multiple immune cell types and stimulates the generation of potentially pathogenic autoantibodies, neutralization of Hb could have beneficial effects.

## Introduction

Systemic Lupus Erythematosus (SLE) is characterized by the presence of antibodies against nuclear, cytoplasmic and cell surface moieties.^1^ Autoimmune hemolytic anemia (AIHA) is observed in SLE patients^2^ and is associated with autoantibodies against red blood cell surface antigens. AIHA can occur in both childhood-onset SLE and adult SLE patients.^3^ Hemolysis results in the release of free hemoglobin (Hb), which normally binds Haptoglobin (Hp) and is cleared via CD163-mediated endocytosis.^4^ In lupus, malaria and leishmaniasis, free Hb concentrations can overwhelm Hp binding capacity, and levels of Hp can be lowered.^5-9^ Hb and heme act as erythrocytic danger-associated molecular patterns which can activate the innate immune system.^10^ Hb-elicited inflammatory responses in endothelial cells^11^, leukocytes^12,13^, and toxicity towards cerebral cortical cells^14^ has been described; Hb can also activate dendritic cells and macrophages^15^, stimulate T cells^16,17^, as well as induce systemic hypertension^18^ and oxidative stress.^19^

Interestingly, patients of lupus, malaria and leishmaniasis harbor anti-Hb autoantibodies in serum, as do aging lupus-prone mice.^20,21^ Immunization of lupus-prone mice with murine Hb results in the generation of antibodies to a wide spectrum of lupus-associated autoantigens and accelerated onset glomerulosclerosis.^21^

In the current study, the inflammatory and immunological properties of ferrous and ferric Hb were explored in detail, based on the premise that biased recognition of the molecule by cells of both the innate and adaptive immune systems may contribute to disease progression in lupus.

## Materials and Methods

### Human samples

This work was carried out in accordance with The Code of Ethics of the World Medical Association (Declaration of Helsinki) for experiments involving humans. In addition, this study was carried out in accordance with the recommendations of the ethical guidelines for biomedical research on human participants laid down by the Indian Council of Medical Research, with informed consent. The protocol was approved by the Institutional Human Ethics Committee of the National Institute of Immunology. SLE patients were diagnosed based on the 2012 Systemic Lupus International Collaborating Clinics (SLICC) classification criteria. Blood was drawn from SLE patients (aged between 21 and 58 years, of North Indian ethnicity) on follow-up, and from ethnicity-, age- and sex-matched healthy donors.

### LPS + R848-induced and Hb-induced cytokine secretion in human peripheral blood mononuclear cells (PBMCs), and preliminary RNA-sequencing

PBMCs isolated from SLE patients and healthy donors were cultured in medium (RPMI-1640 supplemented with 10% fetal bovine serum (FBS; Gibco) and an antibiotic-antimycotic cocktail (Invitrogen), MEM-nonessential amino acids (Gibco), sodium pyruvate (1 mM; Gibco) and 50 μM β-mercaptoethanol (Sigma)), in medium containing human hemoglobin (Hb) (0.5 µM; Sigma Aldrich), or in medium containing a combination of LPS (5 µg/ml; Sigma Aldrich) plus the TLR-7 and TLR-8 agonist Resiquimod (R848, 2 µM, InvivoGen), for 24 h. Supernatants were analyzed for lupus-associated cytokines by ELISA (Invitrogen / R&D). Whether haptoglobin (Hp; Sigma Aldrich) diminished Hb-mediated cytokine release from lupus PBMC was assessed; 0.65 µM Hp (1-1) was incubated with 0.5 µM human Hb (constituting a ratio of Hb:Hp :: 1:1.3) for 2 h at 37°C, and then cultured with PBMCs as described above.

In preliminary RNA-seq analysis, PMBCs (isolated from a healthy donor as well as an SLE patient) were cultured in medium, or in medium containing human hemoglobin (Hb) (0.5 µM; Sigma Aldrich) for 24 h. RNA was isolated using TRIzol (Thermo Fisher Scientific) and subjected to library construction. Transcriptome sequencing was carried out using NOVASeq 6000 (Macrogen Inc, Korea). Reads were mapped to the human genome - Genome Reference Consortium Human Genome Build 38 (GRCh38) using STAR-Aligner; read counts were normalized, and differential expression was calculated using edgeR-Bioconductor package. The Principal Component Analysis (PCA) plot was generated using ggfortify-R CRAN. Up-regulated and down-regulated genes with - FDR < 0.05 with LogFC ≥ 1 or LogFC ≤ −1 - in medium *vs*. Hb-treated PBMCs were represented as volcano plots, generated using the ggplot2-R CRAN package. A pathway and gene ontology enrichment study - with q values less than 0.05 - was carried out using the clusterProfiler Bioconductor package. Kyoto Encyclopedia of Genes and Genomes (KEGG) pathways were viewed using pathview bioconductor package, and heatmaps were generated using the pheatmap - R CRAN package. GEO Accession Numbers: GSE151843 (for healthy donor); GSE141318 (for SLE patient).

### Animals

All animal studies were performed in compliance with the U.S. Department of Health and Human Services Guide for the Care and Use of Laboratory Animals. In addition, this study was carried out in accordance with the recommendations of the Committee for the Purpose of Control and Supervision of Experiments on Animals (CPCSEA), Government of India. The protocol was approved by the institutional animal ethics committee of the National Institute of Immunology. Female, lupus-prone mice NZM2410 (hereafter referred to as NZM) and FVB mice were procured from The Jackson Laboratory, USA.

### Purification of murine Hb and assessment of splenocyte-, B cell-, CD4 T cell-, CD8 T cell- and plasmacytoid dendritic cell-responses to Hb

Murine Hb was isolated from blood derived from NZM mice using standard protocols.^21,22^ Briefly, cold potassium phosphate-buffered saline (PBS; 10 mM, pH 6.5) was added to heparinized blood, and centrifugation carried out at 3,220 g at 4°C for 15 min. After cell lysis, the lysate was dialyzed against PBS and then centrifuged at 1,575 g for 45 min. The supernatant was loaded onto a CM-52 column (Whatman). Eluted fractions, collected upon application of a pH gradient [10 mM potassium phosphate buffer (pH 6.5) and 15 mM potassium phosphate buffer (pH 8.5)], were analyzed by HPLC and SDS-PAGE; electrospray mass spectrometry and N-terminal sequencing were carried out to confirm identity. Data revealed purity of greater than 98%. Ferric (Fe^3+^) Hb was generated by addition of an equimolar concentration of H_2_O_2_ to ferrous (Fe^2+^) Hb; conversion was verified spectrophotometrically (Shimadzu).

B cells, CD4 T cells and CD8 T cells were sorted from splenocytes on FACS Aria (BD Biosciences) using fluorochrome-labeled antibodies (anti-CD19-PECy5, anti-CD4-FITC, anti-CD8-APC; eBiosciences). Plasmacytoid dendritic cells (pDCs) were isolated using a pDC isolation kit (Miltenyi Biotec). Splenocytes, B cells, CD4 T cells, CD8 T cells and pDCs derived from young (2 month-old) and old (8 month-old) FVB and NZM mice were cultured (0.2 x 10^6^ cells/well) in medium, or in medium containing ferrous murine Hb (0.5 µM) or ferric murine Hb (0.5 µM); LPS (0.5 µg/ml; Sigma Aldrich), or LPS (0.5 µg/ml) + ODN 1585 (2 µg/ml; Invitrogen) (the latter for pDCs) were employed as control stimulants. After 24 h, half the medium was replaced, following which cells were incubated for another 24 h. Phorbol myristate acetate (PMA, 50 ng/ml; Sigma Aldrich) and ionomycin (500 ng/ml; Sigma Aldrich) were then added, followed by another incubation for 24 h. Supernatants were assessed for the presence of lupus-associated cytokines by ELISA (Invitrogen / R&D). In analogous cultures of splenocytes, B cells, CD4 T cells, or CD8 T cells, cellular proliferation was determined by assessing the incorporation of ^3^H-Thymidine; LPS (5 µg/ml) or LPS (5 µg/ml) + concanavalin A (5 μg/ml; InvivoGen) were employed as control stimulants for splenocytes and isolated cells respectively.

For the detection of intracellular cytokines in splenocyte cultures, monensin (2 µM; Sigma Aldrich) was added 3 h after the addition of PMA and ionomycin. After 3 h, cells were incubated with Fc block (2 µg, Biolegend). LIVE/DEAD™ fixable violet (Thermo Fisher Scientific) was added, followed by antibody-fluorochrome conjugates (Invitrogen): anti-F4/80-PE-Cy7, anti-CD4-FITC, anti-CD8b-APC and anti-CD19-PE-Cy5. After fixation and permeabilization, fluorochrome-conjugated antibodies to cytokines [anti-IL-6-PE, anti-IL-10-PE, anti-TNF-α-PE, anti-IFN-γ-PE; Invitrogen] were added. After flow cytometry, data was analyzed by FlowJoX software (Tree Star, Inc.).

### Hb-induced signaling

Splenocytes or pDCs isolated from 8 month-old NZM mice were individually incubated with signaling inhibitors [a Stat3 inhibitor (125 nM), staurosporine (PKC inhibitor; 6.25 nM), rapamycin (mTOR inhibitor; 125 nM), H89 (PKA inhibitor; 2 μM), JNK2 (JNK2 inhibitor; 2.5 μM), SB (p38 MAPK inhibitor; 125 nM), PD 98058 (ERK inhibitor; 500 nM), a Stat5 inhibitor (10 μM), Ly (PI3K inhibitor; 1.25 μM)] (Calbiochem) for 1 hr prior to addition of Hb. Supernatants were assessed for the presence of lupus-associated cytokines by ELISA.

### Analysis of Hb-interacting self-moieties

Ferric Hb and ferrous Hb were individually biotinylated using sulfo-NHS-LC-Biotin (Sulfosuccinimidyl-6- {biotin-amido} hexanoate; Thermo Fisher Scientific) and immobilized on streptavidin beads in spin columns (Thermo Fisher Scientific). Lysate derived from Lewis Lung Carcinoma (LLC1) cells (employed as a source of murine “self” moieties) was incubated with Hb beads, and bound moieties were eluted at low pH. Hb binding partners were identified by ESI-LC-MS/MS mass spectrometry (LTQ-Orbitrap Velos, Thermo Fisher Scientific) using Proteome Discoverer 1.3 (Thermo Fisher Scientific). SEQUEST was used for searching protein sequences in murine databases.

### Effect of hemoglobin-apoptotic bleb co-incubation on cytokine and autoantibody responses in murine splenocyte cultures

LLC1 cells were incubated with 40 μM Tamoxifen (Enzo Life Sciences) for 24 h to induce apoptosis, after which apoptotic blebs were isolated. Briefly, cells were centrifuged at 1500 g for 10 min at 25°C; the supernatant was centrifuged at 15700 g for 50 min at 25^°^C. After three washes, the pellet (containing apoptotic blebs) was resuspended in PBS. Apoptotic blebs (5 µg/ml) were incubated with either ferrous Hb (2.5 µM), ferric Hb (2.5 µM), or LPS (5 µg/ml) for 2 h at 37°C. Splenocytes isolated from old NZM and FVB mice were then added, and an incubation carried out for 5 days; cells individually incubated with ferric Hb, ferrous Hb, apoptotic blebs or medium served as negative controls. Supernatants were assessed for the presence of cytokines. Autoreactivity of antibodies in supernatants was assessed by two methods: In the first, moieties in LLC1 cellular lysate were employed as targets on Western blot, employing standard protocols. In the second, reactivity towards lupus-associated autoantigens (dsDNA, Sm, RNP 68K, Ro60, La; Arotec Diagnostics Limited), as well as towards murine ferrous Hb and murine ferric Hb, was assessed by ELISA. Briefly, the autoantigens and murine Hb were individually adsorbed onto ELISA plates (1 μg/well) for 16 h at 4°C; wells were then “blocked” by incubation with 5% BSA in PBS for 2 h at 37°C. Supernatants (diluted 1:5 in 1% BSA in PBS, containing 0.1% Tween 20) were then added, and an incubation carried out for 16 h at 4°C. Following washes with PBST, appropriately diluted goat anti-mouse-HRP antibodies (Jackson ImmunoResearch) were added, and an incubation was carried out for 2 h at 37°C. The plates were washed with PBST, followed by the addition of 50 μl of TMB (Invitrogen). The enzyme reaction was stopped by the addition of 25 μl 2N H_2_SO_4_ (Merck). Absorbance was determined at 450 nm on a BioTek ELISA reader.

### Effect of hemoglobin infusion on cytokine responses, autoantibody responses, and kidney histology

Eight-week-old NZM and FVB mice were intraperitoneally infused with murine ferric Hb (1 mg) or LPS (100 μg) in PBS; respective littermates received PBS as control. Blood samples were collected pre-infusion and at 2 h post-infusion. Cytokines and autoantibodies in plasma were assessed as described above.

The effects of Hb infusion on kidney histology were also assessed. Six-month-old NZM and FVB mice were administered three weekly injections of murine ferric Hb (1 mg); respective littermates received PBS as control. Kidneys were excised and fixed in buffered formalin; 4 μm sections were stained with hematoxylin and eosin. The extent of glomerular damage was assessed by a pathologist in a blinded fashion. Scores were assigned as: Normal (an absence of sclerosis), mild sclerosis (≤ 25% sclerosed glomeruli), moderate sclerosis (26% - 49% sclerosed glomeruli) or severe sclerosis (≥ 50% sclerosed glomeruli).

### Statistical analysis

All data exhibited normal distribution. Results are presented as Mean ± SEM (Standard Error of Mean). The unpaired Student’s t-test, one-way ANOVA and two-way ANOVA were used to calculate statistical significance, as appropriate. P values of less than 0.05 were considered significant.

## Results

### LPS + R848-induced and human Hb-induced cytokine responses in human PBMCs

PBMCs isolated from healthy individuals and SLE patients secreted equivalent levels of IL-6, IL-10 and TNF-α in response to LPS + R848. In contrast, PBMCs isolated from SLE patients secreted enhanced levels of these cytokines in response to human Hb, compared to PBMCs derived from healthy donors (Figure 1 A-C). In PBMCs isolated from SLE patients, secretion of Hb-induced IL-10 was completely abrogated, and secretion of IL-6 and TNF-α decreased to a significant extent, upon the addition of Hp (Figure 1 D).

**Figure 1.**
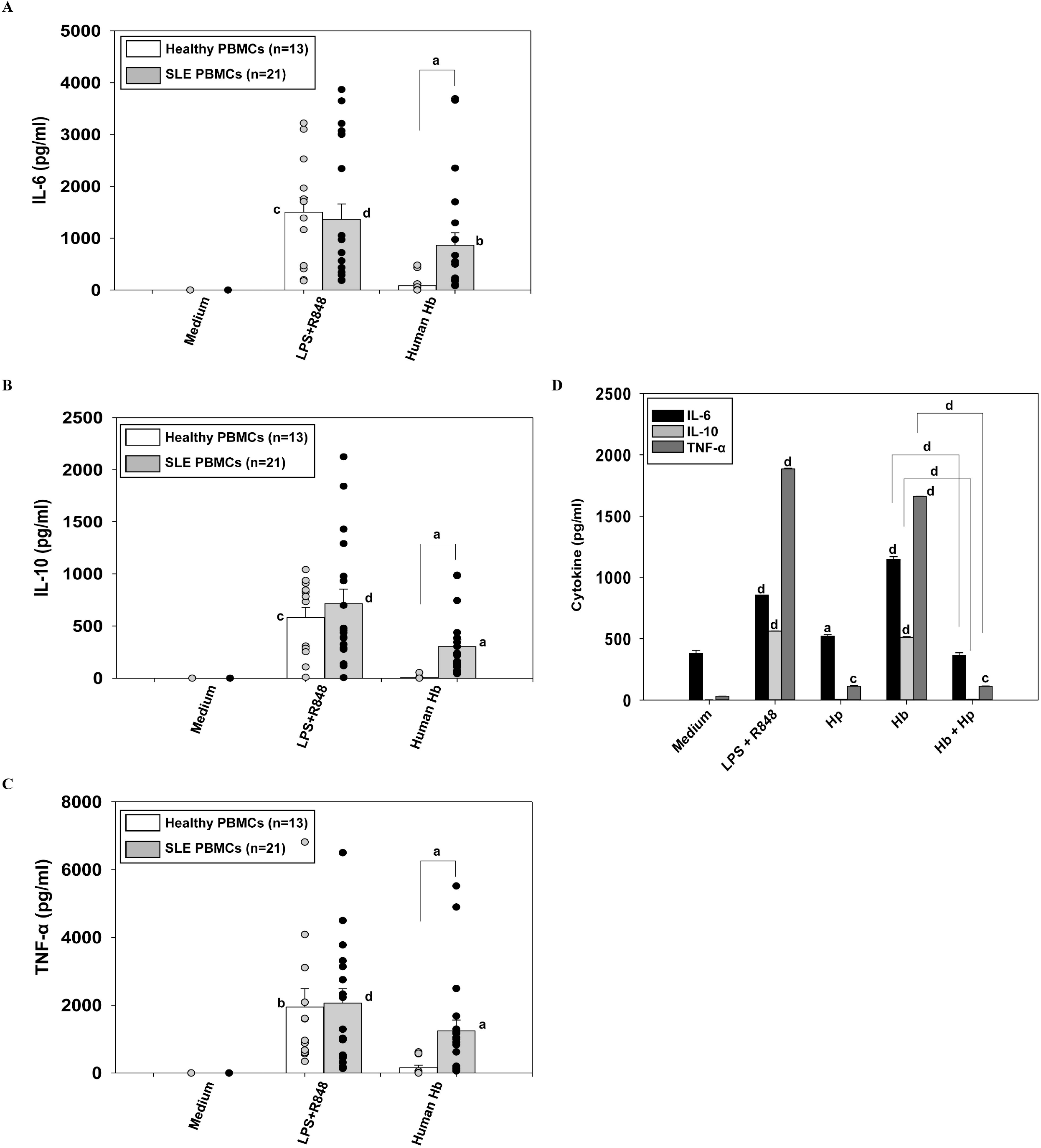
LPS + R848-induced and human Hb-induced cytokine secretion from human PBMCs. Human PBMCs isolated from healthy subjects (n=13) or SLE patients (n=21) were incubated with medium, or with medium supplemented with 5 µg/ml LPS + 2 µM R848, or with 0.5 µM human Hb. (A-C) Levels of (A) IL-6, (B) IL-10 and (C) TNF-α in supernatants. Dots represent responses of individual subjects/patients. (D) IL-6, IL-10 and TNF-α were estimated in supernatants of SLE PBMCs incubated with medium, LPS + R848, Hp, human Hb, or human Hb + Hp. Data represent Mean ± SEM. ^a^p<0.05, ^b^p<0.01, ^c^p<0.001, ^d^p<0.0001 vs medium, unless otherwise indicated.

### LPS-induced and murine Hb-induced splenocyte cytokine and proliferative responses in murine cells

As expected, LPS was generally inflammatory when incubated with splenocytes isolated from (young and old) healthy and lupus-prone mice. In contrast, splenocytes derived from aging NZM mice released higher levels of IFN-γ, IL-6, IL-8, IL-10 and TNF-α upon incubation with murine ferrous Hb or murine ferric Hb, than did splenocytes from young NZM mice, or from old FVB mice; murine ferric Hb elicited significantly greater responses than ferrous Hb (Figure 2 A-E). The presence of intracellular cytokines was also assessed in NZM splenocyte cultures; while representative gating strategies, as well as representative assessments of antibody specificity on different splenocyte subpopulations, are depicted in Supplemental Figure 1, data on all splenocytes was acquired and analyzed for the purposes of this study. Murine ferrous Hb and murine ferric Hb induced the intracellular expression of IFN-γ, IL-6, IL-10 and TNF-α in B cells, monocytes, CD4 T cells and CD8 T cells; a higher number of B cells, monocytes and CD4 T cells demonstrated the presence of the cytokines when incubated with murine ferric Hb than when incubated with murine ferrous Hb (Figures 2 F-I).

**Figure 2.**
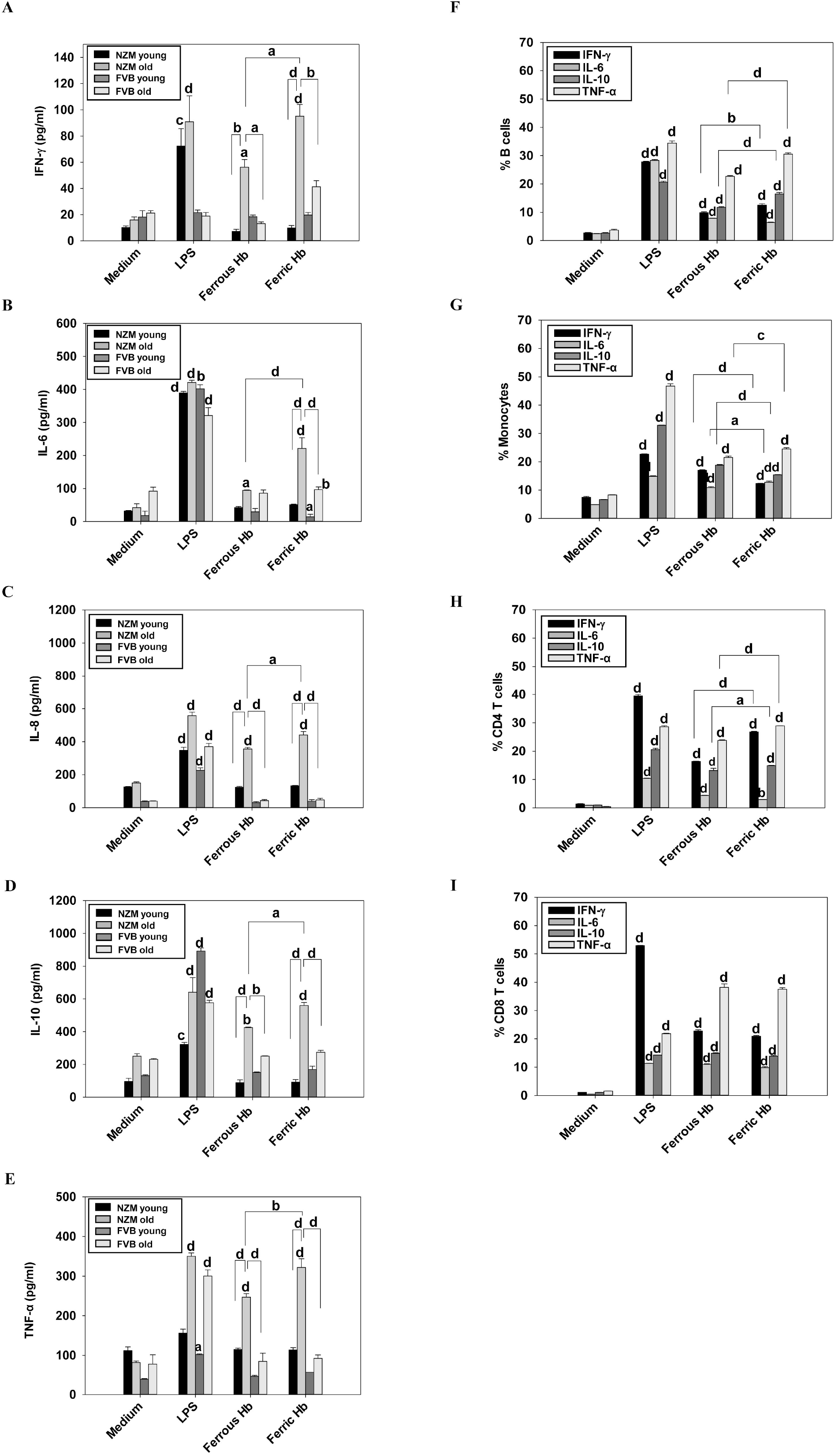
LPS-induced and murine Hb-induced cytokine secretion from splenocytes isolated from NZM and FVB mice. Splenocytes from young (2 month-old) and old (8 month-old) NZM mice (n=4) and FVB mice (n=4) were incubated with medium, or with medium supplemented with 5 µg/ml LPS, 0.5 µM murine ferrous Hb, or 0.5 µM murine ferric Hb. (A-E) Levels of (A) IFN-γ, (B) IL-6, (C) IL-8, (D) IL-10 and (E) TNF-α in supernatants. (F-I) Splenocytes from old (8 month-old) NZM mice were incubated with medium, or with medium supplemented with 5 µg/ml LPS, 0.5 µM murine ferrous Hb, or 0.5 µM murine ferric Hb. Intra-cellular IFN-γ, IL-6, IL-10 and TNF-α was assessed in (F) B cells, (G) Monocytes, (H) CD4 T cells and (I) CD8 T cells. Each experiment was carried out four times. Data represent Mean ± SEM. ^a^p<0.05, ^b^p<0.01, ^c^p<0.001, ^d^p<0.0001 vs medium, unless otherwise indicated.

In general, the trends observed for splenocytes held true for isolated cells as well. Murine ferrous Hb and murine ferric Hb led to higher release of lupus-associated cytokines from B cells (IFN-γ, IL-6, IL-10; Figure 3 A-C) and CD4 T cells (IFN-γ, TNF-α, IL-10; Figure 3 D-F) isolated from old NZM mice, compared with respective cells isolated from young NZM mice, or old FVB mice. CD8 T cells isolated from old NZM mice were also more responsive to murine ferric Hb than cells isolated from young NZM mice or from old FVB mice (data not shown).

**Figure 3.**
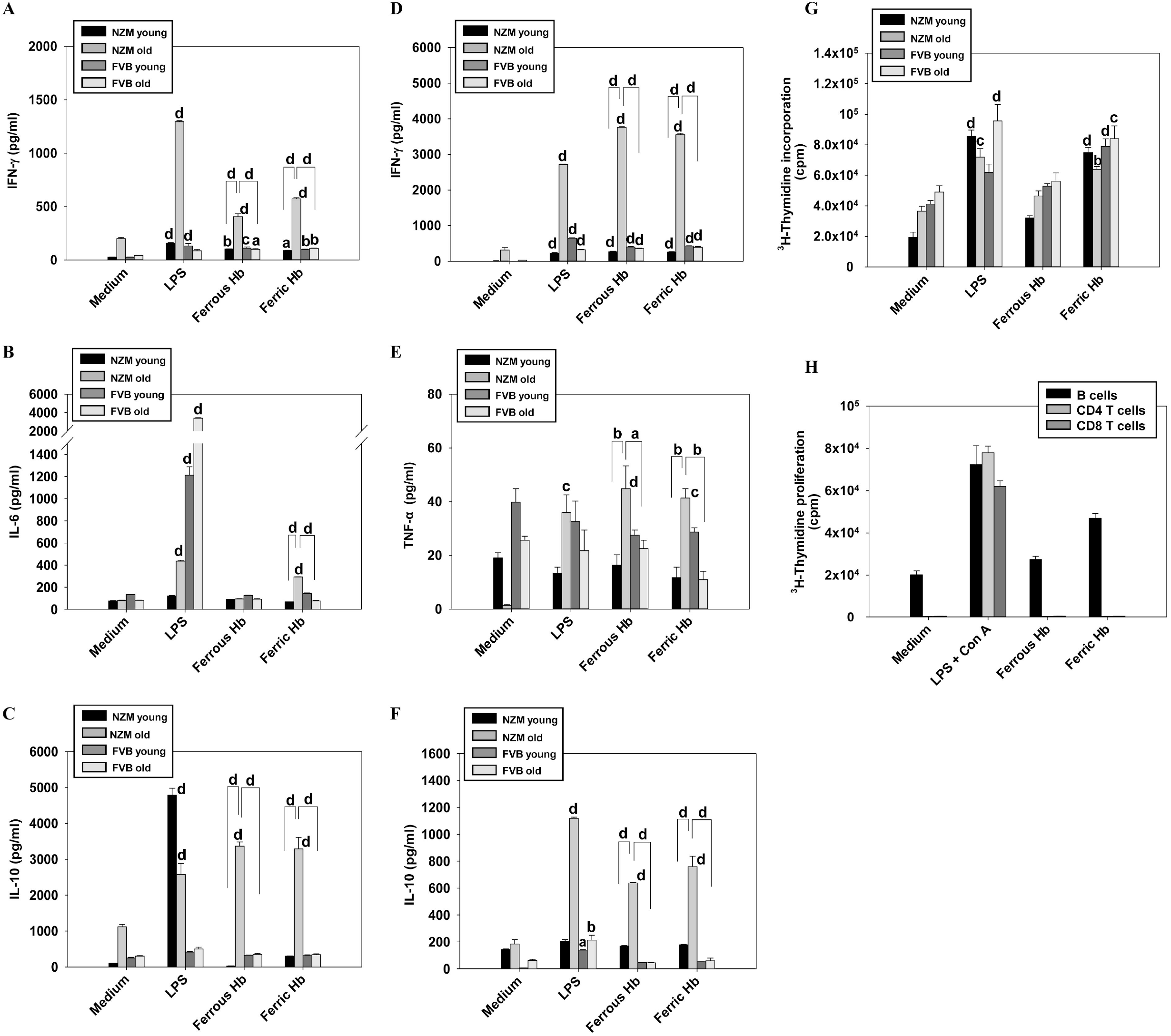
LPS-induced and murine Hb-induced cytokine and proliferative responses of lymphocytes isolated from NZM and FVB mice. (A-F) Levels of cytokines (as indicated) in the supernatants of (A-C) B cells and (D-F) CD4 T cells (isolated from young (2 month-old) and old (8 month-old) NZM mice (n=4) and FVB mice (n=4)) incubated with medium, or with medium supplemented with 5 µg/ml LPS, 0.5 µM murine ferrous Hb, or 0.5 µM murine ferric Hb. (G) Proliferative responses of splenocytes isolated from young (2 month-old) and old (8 month-old) NZM mice (n=4) and FVB mice (n=4), cultured as described above. (H) Proliferative responses of B cells, CD4 T cells or CD8 T cells isolated from old NZM mice (n=4), incubated with medium, or with medium supplemented with 5 µg/ml LPS + 5 μg/ml Concanavalin A (ConA), 0.5 µM murine ferrous Hb, or 0.5 µM murine ferric Hb. Each experiment was carried out four times. Data represent Mean ± SEM. ^a^p<0.05, ^b^p<0.01, ^c^p<0.001, ^d^p<0.0001 vs medium, unless otherwise indicated.

While murine ferric Hb induced proliferative responses in splenocytes derived from both NZM and FVB mice (irrespective of age), murine ferrous Hb did not (Figure 3 G); proliferation was preferentially observed in isolated B cells, as opposed to isolated CD4 T cells or CD8 T cells (Figure 3 H).

### LPS-induced and murine Hb-induced pDC cytokine secretion

While LPS was generally inflammatory when incubated with pDCs derived from both NZM and FVB mice, both murine ferrous Hb and ferric Hb induced heightened secretion of several inflammatory cytokines (IFN-γ, IL-6, IL-8, IL-10, IL-12, TNF-α) from pDCs derived from old NZM mice, compared with cells derived from young NZM mice or old FVB mice; ferric Hb invariably induced higher cytokine responses than did ferrous Hb (Figure 4 A-F).

**Figure 4.**
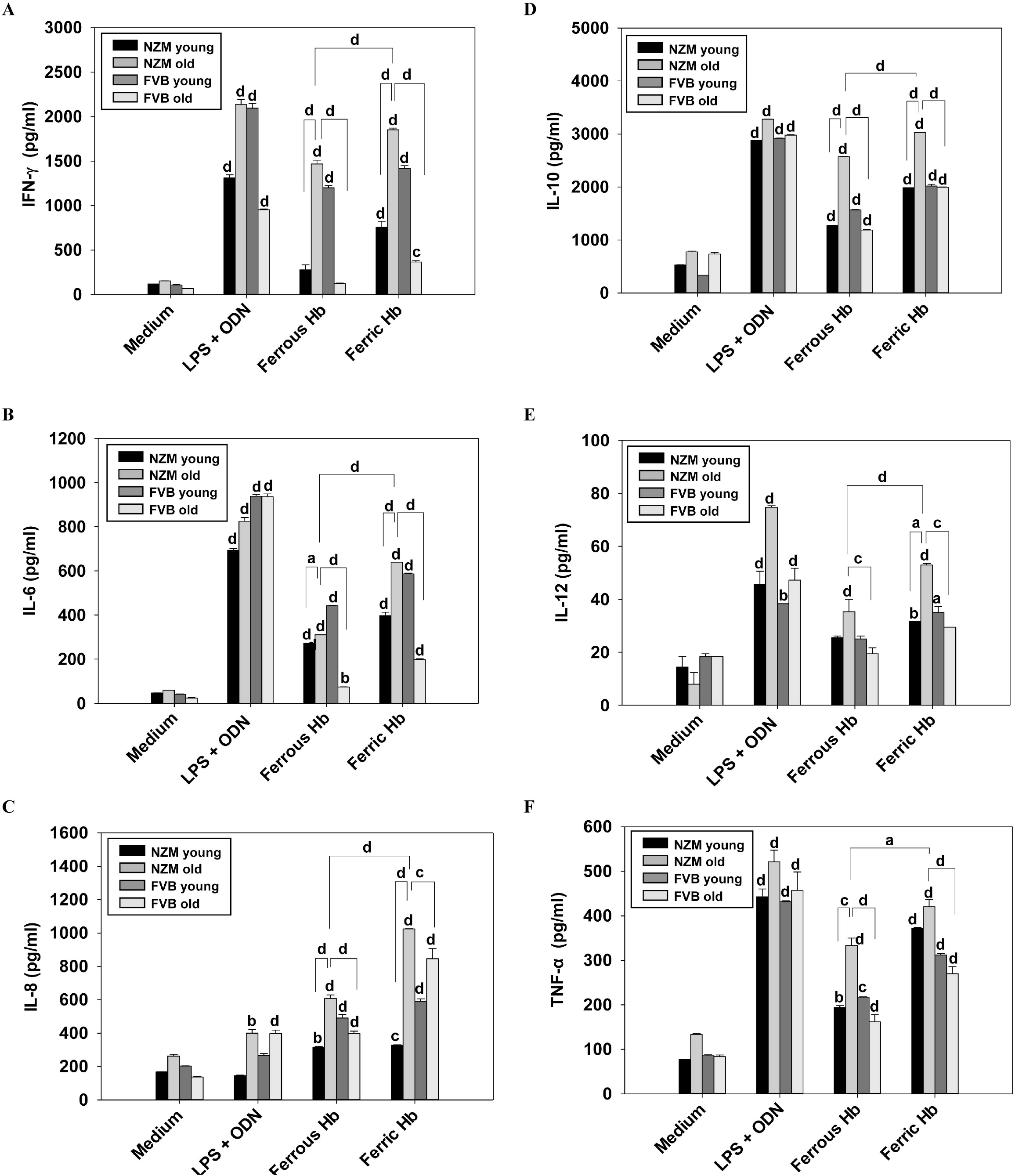
LPS+ODN 1584-induced and murine Hb-induced cytokine secretion from pDCs isolated from NZM and FVB mice. pDCs from young (2 month-old) and old (8 month-old) NZM mice (n=4) and FVB mice (n=4) were incubated with medium, or with medium supplemented with 5 µg/ml LPS + 2 µM ODN 1585, 0.5 µM murine ferrous Hb, or 0.5 µM murine ferric Hb. Levels of (A) IFN-γ, (B) IL-6, (C) IL-8, (D) IL-10, (E) IL-12 and (F) TNF-α in supernatants. Each experiment was carried out three times. Data represent Mean ± SEM. ^a^p<0.05, ^b^p<0.01, ^c^p<0.001, ^d^p<0.0001 vs medium, unless otherwise indicated.

### Murine Hb-induced signaling in splenocytes and pDCs

Splenocytes and pDCs isolated from old NZM mice were incubated with murine ferric Hb in the presence of individual signaling inhibitors; effects on IL-6, IL-10, TNF-α and IL-8 secretion were assessed. In splenocytes incubated with murine ferric Hb, addition of a Stat3 inhibitor diminished the secretion of IL-6 and TNF-α. Staurosporine affected the release of IL-6 and almost completely abrogated the release of IL-10 and TNF-α. Rapamycin significantly decreased the release of IL-6, IL-10 and TNF-α, and completely ablated IL-8 release. The addition of H89 led to the decreased production of IL-6, IL-10 and TNF-α. The JNK2 inhibitor caused the decreased release of IL-6 and IL-10, while completely halting the release of IL-8. The addition of SB greatly decreased IL-8, while diminishing IL-6, IL-10 and TNF-α secretion to a lesser extent (Supplemental Figure 2A). In pDCs incubated with murine ferric Hb, the Stat3 inhibitor led to a decrease in the release of TNF-α and IL-8. Staurosporine, while affecting the release of TNF-α, completely ablated IL-8 secretion. Rapamycin diminished the release of IL-6 and abolished the secretion of IL-8. H89 attenuated the release of IL-10 and completely inhibited IL-8 secretion. The JNK2 inhibitor prevented the secretion of IL-8, while SB decreased TNF-α release while completely preventing IL-8 release (Supplemental Figure 2B). The down-modulatory effects of the inhibitors therefore varied, depending on the cytokine readout as well as the cell type. The potential significance of these observations is discussed below.

### Enumeration of Hb-autoantigen interaction

Pull-downs were carried out to assess murine Hb-self moiety interaction; Supplemental Figure 3A depicts SDS-PAGE of eluted fractions. Moieties binding control (uncoupled) beads were excluded from analyses. Two hundred and eighty-four self-moieties were pulled down by murine ferrous Hb beads and 170 self-moieties by murine ferric Hb beads (Supplemental Figure 3B). Of these, known lupus-associated autoantigens were short-listed; 24 lupus-associated autoantigens were found to bind both murine ferrous Hb and murine ferric Hb, 13 bound only murine ferrous Hb, and 5 bound only murine ferric Hb (Supplemental Table I). In other words, of the 284 self-moieties pulled down with murine ferrous Hb, 37 (13.02%) were identified as lupus-associated autoantigens, and of the 170 self-moieties pulled down with murine ferric Hb, 29 (17.02%) were identified as lupus-associated autoantigens.

### The effect of apoptotic blebs on Hb-induced cytokine and autoantibody responses in vitro

Given current and previous^21^ evidence of the ability of Hb to interact with autoantigens, several of which are found in apoptotic blebs, whether Hb-apoptotic bleb co-incubation acted as a positive stimulus for cytokine and autoantibody secretion was assessed.

#### Cytokines

The combination of murine ferric Hb + apoptotic blebs (as opposed to murine ferrous Hb + apoptotic blebs) led to the enhanced secretion of lupus-associated cytokines (IFN-γ, IL-6, IL-8, IL-10, TNF-α) when incubated with NZM (but not FVB) splenocytes, compared to when individual components were employed. Responses to LPS + apoptotic blebs were less discriminatory between the two murine strains (Figure 5 A-E).

**Figure 5.**
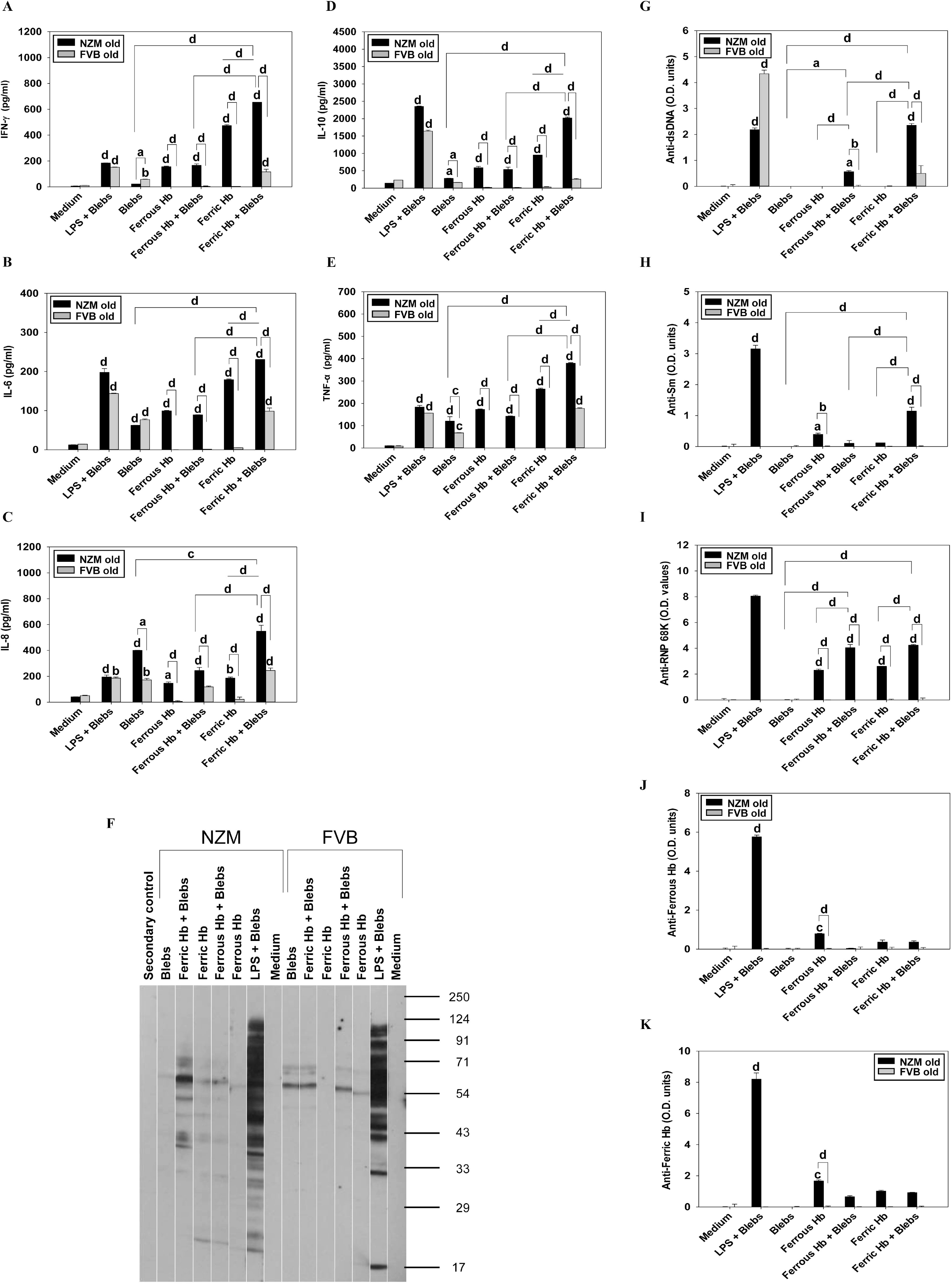
Effect of murine Hb-apoptotic bleb co-incubation on the secretion of cytokines and autoantibodies from splenocytes isolated from NZM and FVB mice. Splenocytes isolated from old (8 month-old) NZM mice (n=4) and FVB mice (n=4) were incubated with medium, or with medium supplemented with 5 µg/ml LPS + 5 µg/ml apoptotic blebs, apoptotic blebs, 2.5 µM murine ferrous Hb, 2.5 µM murine ferrous Hb + 5 µg/ml apoptotic blebs, 2.5 µM murine ferric Hb, or 2.5 µM murine ferric Hb + 5 µg/ml apoptotic blebs. (A-E) Levels of (A) IFN-γ, (B) IL-6, (C) IL-8, (D) IL-10, and (E) TNF-α in supernatants. (F) Western blot on LLC1 lysate, depicting autoantibody reactivity in supernatants. “Secondary control” refers to a strip incubated with just the secondary antibody. (G-K) ELISAs depicting reactivity towards (G) dsDNA, (H) Sm, (I) RNP 68K, (J) Ferrous Hb, and (K) Ferric Hb. Each experiment was carried out four times. Data are expressed as O.D. Units (O.D. x dilution factor) and represent Mean ± SEM. ^a^p<0.05, ^b^p<0.01, ^c^p<0.001, ^d^p<0.0001 vs medium, unless otherwise indicated.

#### Autoantibodies

Incubation of splenocytes derived from NZM mice with murine ferric Hb + apoptotic blebs (as opposed to murine ferrous Hb + apoptotic blebs) led to substantial increases in autoantibodies levels in culture supernatants, compared to when individual components were employed. On Western blot, a diversification of secreted autoantibody specificity was evident when both moieties were present in culture; several additional self-moieties were targeted. Equivalent diversity in autoantibody specificity was not observed in culture supernatants when splenocytes derived from FVB mice were incubated with murine ferric Hb + apoptotic blebs (Figure 5 F). NZM splenocytes, upon incubation with murine ferric Hb + apoptotic blebs, secreted antibodies reactive to the lupus-associated autoantigens dsDNA, Sm and RNP 68K; incubation with murine ferrous Hb + apoptotic blebs resulted in significantly lower levels of such antibodies in supernatants. Incubation of splenocytes derived from FVB mice with murine ferric Hb + apoptotic blebs did not result in the heightened secretion of antibodies against these autoantigens (Figure 5 G-I). Interestingly, antibodies reactive to ferrous Hb or ferric Hb were not enhanced in supernatants derived from these cultures (Figure 5 J, K).

### Enumeration of Hb-induced cytokine and autoantibody responses in vivo

#### Cytokines

Infusion of murine ferric Hb into NZM mice induced higher levels of several lupus-associated cytokines (IFN-γ, IL-6, IL-8, IL-10, TNF-α) than did infusion into FVB mice; similar bias between the two murine strains was not apparent upon the infusion of LPS (Figure 6 A-E).

**Figure 6.**
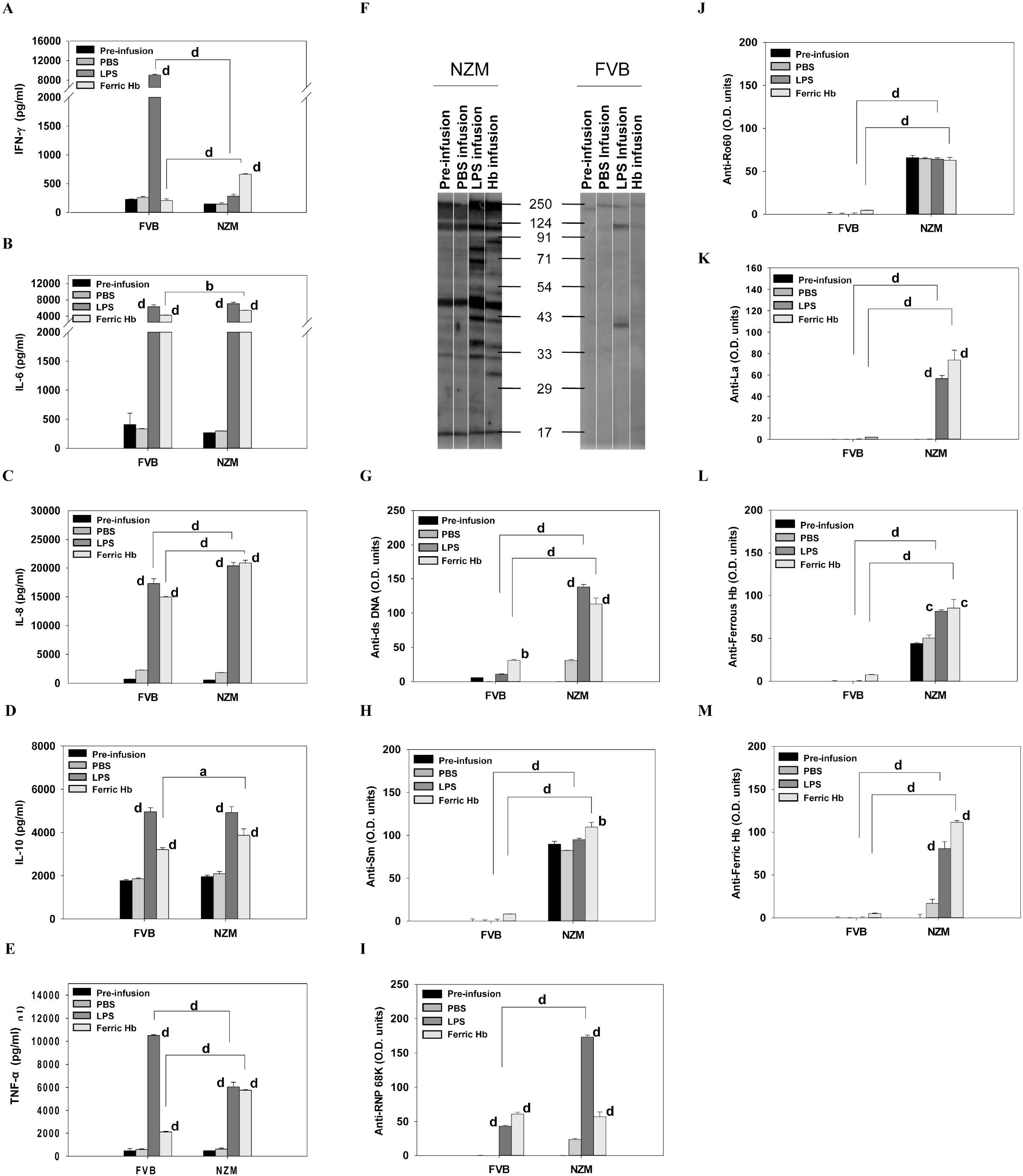
Murine Hb-induced cytokine and autoantibody secretion in NZM and FVB mice *in vivo*. Eight-week-old NZM mice (n=8) and FVB mice (n=8) were infused with PBS, or with PBS containing LPS (100 μg) or murine ferric Hb (1 mg). (A-E) Levels of (A) IFN-γ, (B) IL-6, (C) IL-8, (D) IL-10, and (E) TNF-α in plasma pre-infusion and 2 h post-infusion. (F) Western blot on LLC1 lysate, depicting autoantibody reactivity in plasma pre-infusion and 2 h post-infusion. (G-M) ELISAs depicting reactivity of antibodies in plasma (pre-infusion and post-infusion) towards (G) dsDNA, (H) Sm, (I) RNP 68K, (J) Ro60, (K) La, (L) Ferrous Hb, and (M) Ferric Hb. Data are expressed as O.D. Units (O.D. x dilution factor) and represent Mean ± SEM. ^a^p<0.05, ^b^p<0.01, ^c^p<0.001, ^d^p<0.0001 vs pre-infusion, unless otherwise indicated.

#### Autoantibodies

As expected, the infusion of PBS did not enhance autoreactivity (over respective pre-infusion levels) in either murine strain, as assessed by Western blot. Infusion of LPS induced the generation of autoantibodies in NZM mice and (to a lesser extent) in FVB mice. Significantly, the infusion of murine ferric Hb into NZM mice (but not into FVB mice) led to increased plasma autoantibody levels; several additional moieties on cellular lysate were targeted, compared with autoantibodies in plasma derived pre-infusion and post-PBS infusion (Figure 6 F). Heightened autoreactivity towards the autoantigens dsDNA, Sm, RNP 68K and La was observed; reactivity to Ro60 was not enhanced, presumably due to high pre-existing autoantibody levels. With the exception of RNP 68K, the infusion of murine ferric Hb into FVB mice generated significantly lower levels of autoantibodies against the autoantigens (Figure 6 G-K). Of additional interest was the fact that infusion of murine ferric Hb in NZM mice (but not in FVB mice) induced the generation of antibodies towards murine ferrous Hb (Figure 6 L) and murine ferric Hb (Figure 6 M), further indication of the auto-immunogenic nature of Hb specifically in a lupus milieu.

#### Kidney histology

Histological analysis revealed that infusion of murine ferric Hb in NZM mice induced glomerulosclerosis, increases in the mesangial matrix, and a thickening of the basement membrane; similar increases in abnormalities were not observed in NZM mice infused with PBS. Infusion of murine Hb in FVB mice did not induce changes similar to those induced in NZM mice (Figure 7 A). Quantification of the data revealed a significant increase in the number of severely sclerotic glomeruli in Hb-infused NZM mice, compared with PBS-infused NZM mice and with Hb-infused FVB mice (Figure 7 B).

**Figure 7.**
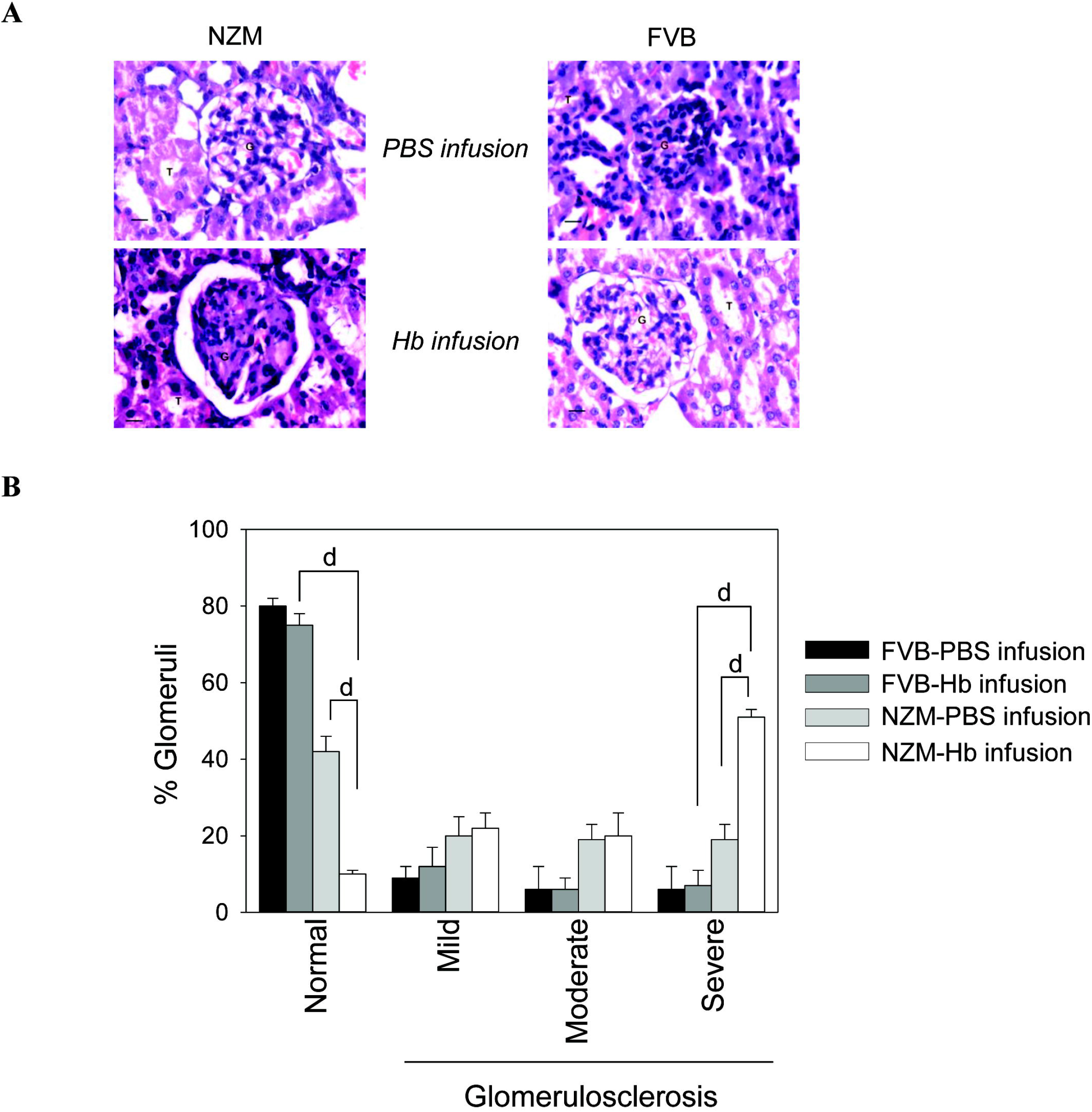
Effect of murine Hb infusion on the kidneys in NZM and FVB mice. Six-month-old NZM mice (n=6) and FVB (n=6) mice were infused (once a week, for three weeks) with PBS or with PBS containing murine ferric Hb (1 mg). (A) Histology of hematoxylin-eosin stained kidney sections post-infusion. (B) Quantification of pathological changes in kidney glomeruli post-infusion. Scores were assigned as: Normal (an absence of sclerosis), Mild Sclerosis (≤ 25% sclerosed glomeruli), Moderate Sclerosis (26% - 49% sclerosed glomeruli) or Severe Sclerosis (≥ 50% sclerosed glomeruli). Data represent Mean ± SEM. ^d^p<0.0001.

## Discussion

In diseases characterized by hemolysis like lupus, malaria and leishmaniasis, free Hb can overwhelm clearance mechanisms.^5-9^ Oxidation of free Hb to its ferric and ferryl forms can result, as can the subsequent release of heme.^23^ The inflammatory and pathological effects of Hb and heme on cells of both immune and non-immune lineage have been well-documented.^10-15, 18, 19, 24-26^

Emerging evidence describes the antigenicity of Hb in lupus patients; antibodies reactive to human Hb have been reported in the sera of malaria and leishmaniasis patients as well^20,21^, but not in the sera of patients of autoimmune and inflammatory disorders not associated with hemolysis.^21^ Higher circulating levels of free murine Hb have been documented in lupus-prone mice, and antibodies to Hb arise in such mice as they age; adherence of anti-Hb antibodies to the lungs and kidneys in these mice is suggestive of a pathological role.^20,21^ Immunization of lupus-prone mice with murine Hb results in the generation of antibodies to a wide spectrum of lupus-associated autoantigens and the accelerated onset of glomerulosclerosis.^21^ These studies strongly suggest that there exists differential immune recognition of Hb in a milieu characterized by systemic autoimmune aberration, a contention supported by the fact that lupus-prone mice express higher anti-Hb B cell precursor frequencies.^21^

Here, the inflammatory and immunogenic attributes of Hb were delineated in some detail, in the context of human and murine lupus. Post Hb-incubation, PBMCs from SLE patients generated higher levels of IL-6, IL-10 and TNF-α than did PBMCs derived from healthy donors, even though responses to a polyclonal stimulus were similar. These cytokines are of relevance to different aspects of lupus pathology. Higher IL-6 levels occur in SLE patients during active hematological disease,^27,28^ and IL-6 and IL-10 act to increase the secretion of erythrocyte-reactive autoantibodies from PBMCs derived from AIHA patients.^29^ BT063 (an anti-IL10 mAb) and Vobarilizumab (an anti-IL-6 receptor nanoantibody) have been investigated as potential therapeutics in SLE patients.^30,31^ Further, serum levels of TNF-α exhibit correlation with disease activity in SLE patients.^32^

The triple requirements for robust inflammatory effects of Hb on murine cells (belonging to three lineages of the adaptive immune system and two lineages of the innate immune system) were delineated in this study: a) the preference for a lupus genotype; b) a preference for advanced age, and c) a preference, in most instances, for murine ferric Hb over murine ferrous Hb. Some insights of direct relevance to lupus pathology are discussed here.

While evidence indicates the enhanced phosphorylation of ERK, STAT3 and JNK in NZM splenocytes (versus FVB splenocytes) upon incubation with murine ferric Hb (data not shown), inhibitor studies have revealed additional aspects of Hb-mediated signaling in NZM splenocytes; particular signaling inhibitors exert inhibitory effects towards the secretion of particular cytokines. Comparison between signaling events in splenocytes versus pDCs revealed an additional level of complexity in murine Hb action. Some examples: The STAT3 inhibitor significantly decreased IL-6 production in murine Hb-stimulated splenocytes but had no effect on IL-6 release by pDCs. Additionally, PKC inhibition (by Staurosporine) ablated IL-8 release in murine Hb-stimulated pDCs but had no effect on IL-8 release in murine Hb-stimulated splenocytes. Use of PD 98058 (an ERK inhibitor), a Stat5 inhibitor, and Ly (a PI3K inhibitor) essentially confirmed observations of differential inhibitory effects on splenocytes, depending on the cytokine readout (data not shown).

These results strongly suggest that murine Hb triggers multiple signaling pathways in NZM mice, and that such events can be cell-dependent. LPS is known to stimulate multiple signaling pathways in other systems^33-37^, and data suggests that it triggers several pathways in NZM splenocytes as well; though, as with Hb, particular pathways appeared responsible for the secretion of specific cytokines, several differences were observed in terms of the relative efficacy of specific inhibitors to inhibit specific cytokines (data not shown). The implications of the unique inflammatory properties of Hb in the context of lupus are under study.

Hb is known to demonstrate an inflammatory synergy with some exogenous TLR ligands (such as LPS and LTA^38^); at least in some instances, this effect has been attributed to physical association. Since several ribonucleoprotein autoantigens, as well as dsDNA, that are extruded into apoptotic blebs as cells die also act as endogenous TLR ligands^39,40^, whether Hb-autoantigen complexes play a role in lupus is interesting to consider. Another reason to investigate Hb-autoantigen interactions is the potential impact such associations may have on epitope spreading, the time-dependent diversification of autoantigenic targets that is characteristic of lupus.^41,42^ For physically-associated antigens, a break in T cell tolerance to one component is believed to enable activated T cells to “help” B cells (specific for other component(s) which internalize the complex), resulting in diversification of the autoantibody repertoire. Previous work has determined that Hb and/or heme can interact with several lupus-associated autoantigens - ATP synthase subunit beta, fructose-bisphosphate aldolase A, peroxiredoxin-1, peroxiredoxin-2^43^, apolipoprotein A-I^44^, complement C4-B^45^, superoxide dismutase [Cu-Zn], carbonic anhydrase, actin, and band 3^46^. Work in our lab has previously identified additional autoantigens (La, Sm, Ro52, Ro60, RNP 68K) as being Hb interactors.^21^ Using an unbiased Hb pull-down approach, the current work led to the enumeration of more Hb-interacting autoantigens, including Sm (D3, D2), calreticulin, 60S ribosomal P2, annexin (A4, A11), HMGB1, histone-1, RNA polymerase (I, II, III), as well as many hnRNPs and snRNPs. While the potential biological roles of individual Hb-autoantigen complexes await future elucidation, co-incubation of ferric (but not ferrous) murine Hb and apoptotic blebs (which contain packaged autoantigens) with NZM (but not FVB) splenocytes resulted in inflammatory and immunogenic synergy; heightened levels of lupus-associated cytokines were generated, and greater secretion of autoantibodies, including of specificities associated with pathological sequelae in lupus, was observed. These results are strongly suggestive of a disease-influencing role of Hb-autoantigen complexes in lupus.

Immunization of NZM mice with murine Hb was previously shown to lead to epitope spreading to other lupus-associated autoantigens^21^; those studies employed a strong adjuvant. In current work, simple infusion of murine Hb into lupus-prone NZM mice resulted in the generation of higher levels of several lupus-associated inflammatory cytokines than did infusion into healthy mice; in contrast, and in keeping with previous reports^47^, infusion of LPS into healthy mice elicited higher levels of some cytokines, findings again attesting to differences in downstream events triggered by the two inflammatory moieties in the two murine strains. Infusion of Hb into NZM mice also led to the enhanced generation of autoantibodies (including against dsDNA), and resulted in glomerulosclerosis; whether these effects are a consequence of the ability of Hb to interact with DNA-binding proteins (HMGB1, histone-1 and RNA polymerase), as this study has shown, remains to be determined.

Preliminary RNA-seq analysis, on PBMCs isolated from an SLE patient and a healthy donor, revealed that the former responded to Hb in an enhanced fashion (Supplemental Figure 4A). Expectedly, mRNAs for several proteins involved in the TNF signaling, as well as in JAK-STAT signaling (including for IL-6 and IL-10), were enhanced (Figure Supplemental Figure 4B). Amongst other effects, mRNAs for IL-1β, oncostatin M (OSM), CXCL1, CXCL5 and IL-2RA were up-modulated. IL-1β deficient mice are resistant to the development of experimental lupus^48^, and levels of OSM are associated with SLE disease activity in humans.^49^ Increased glomerular expression of CXCL1 has been documented on renal biopsies of lupus nephritis patients^50^, and CXCL5 causes neutrophil recruitment, contributing to Th17-driven glomerulonephritis.^51^ Enhanced transcription of IL-2RA (CD25) is observed in activated CD4 T cells, and studies suggest that serum levels of the molecule can serve as a marker of disease activity.^52^ Though these initial leads are interesting, validation on a larger number of subjects is obviously critical.

In conclusion, this work describes the unique inflammatory and immunologic stimulus Hb provides in a lupus milieu. Along with previous studies, it further strengthens the case for the classification of Hb as a *bona fide* autoantigen. Interestingly, Hb was incapable of stimulating the release of IFN-α (believed to be critical for the initiation of lupus^53^), either *in vitro* or *in vivo* (data not shown), suggesting that the inflammatory stimulus Hb specifically mediates may contribute to disease progression, but not necessarily to disease initiation. Neutralizing the inflammatory and immunogenic actions of Hb in lupus (for example, by the use of Hp, which is increasingly considered a therapeutic option in humans for the amelioration of Hb-driven inflammation and thrombosis^54^) may prove clinically beneficial.

## Acknowledgements

The authors thank Dr. Ruchi Sachdeva and Dr. Monika Malik for critical scientific inputs.

## Author contributions

HS, UK and RP conceptualized the study and decided on methodology, RP was responsible for funding acquisition and supervised the study, HS and AB carried out the investigations, HS and RP were involved in data curation and formal analysis. HS, AB and RP analyzed the data. HS wrote the original draft of the manuscript. RP reviewed and edited the manuscript.

