## Supplemental Table for "Hemoglobin drives inflammation and initiates antigen spread and nephritis in lupus"

| **Accession** | **Description** | **Score**  **Ferrous Hb** | **Coverage**  **Ferrous Hb** | **Score Ferric Hb** | **Coverage Ferric**  **Hb** | **MW [kDa]** |
| --- | --- | --- | --- | --- | --- | --- |
| P16045 | Galectin-1 | 42.55 | 80.74 | 15.12 | 37.78 | 14.9 |
| P99027 | 60S acidic ribosomal protein P2 | 22.97 | 69.57 | 8.42 | 26.96 | 11.6 |
| P68369 | Tubulin alpha-1A chain | 109.79 | 50.33 | 96.83 | 49.00 | 50.1 |
| Q61171 | Peroxiredoxin-2 | 17.87 | 43.43 | 8.73 | 21.72 | 21.8 |
| Q9R0P3 | S-formylglutathione hydrolase | 25.91 | 42.55 | 12.74 | 14.18 | 31.3 |
| P63323 | 40S ribosomal protein S12 | 10.75 | 34.85 | 7.66 | 34.85 | 14.5 |
| P63101 | 14-3-3 protein zeta/delta | 35.13 | 35.10 | 26.37 | 35.10 | 27.8 |
| P63158 | High mobility group protein B1 | 24.32 | 27.44 | 14.98 | 19.07 | 24.9 |
| P61979 | Heterogeneous nuclear ribonucleoprotein K | 34.88 | 29.37 | 22.13 | 17.28 | 50.9 |
| P62317 | Small nuclear ribonucleoprotein Sm D2 | 7.46 | 32.20 | 14.35 | 16.10 | 13.5 |
| P17095 | High mobility group protein HMG-I/HMG-Y | 6.76 | 23.36 | 11.64 | 22.43 | 11.6 |
| P62849-2 | Isoform 2 of 40S ribosomal protein S24 | 6.13 | 20.77 | 5.91 | 29.23 | 15.1 |
| Q9CX86 | Heterogeneous nuclear ribonucleoprotein A0 | 17.82 | 23.61 | 11.89 | 16.72 | 30.5 |
| P49312 | Heterogeneous nuclear ribonucleoprotein A1 | 14.39 | 22.19 | 8.11 | 13.75 | 34.2 |
| O88569-3 | Isoform 3 of Heterogeneous nuclear ribonucleoproteins A2/B1 | 9.83 | 18.94 | 8.37 | 11.63 | 32.4 |
| P62320 | Small nuclear ribonucleoprotein Sm D3 | 17.21 | 23.81 | 8.51 | 23.81 | 13.9 |
| Q99020 | Heterogeneous nuclear ribonucleoprotein A/B | 36.65 | 23.16 | 22.38 | 17.89 | 30.8 |
| Q8C1B7-3 | Isoform 3 of Septin-11 | 35.15 | 16.00 | 38.26 | 19.06 | 48.9 |
| P43275 | Histone H1.1 | 11.81 | 18.78 | 12.61 | 11.74 | 21.8 |
| P47962 | 60S ribosomal protein L5 | 9.23 | 14.14 | 15.94 | 16.50 | 34.4 |
| P32067 | Lupus La protein homolog | 11.25 | 10.60 | 14.14 | 12.53 | 47.7 |
| P26041 | Moesin | 23.77 | 14.56 | 4.46 | 3.29 | 67.7 |
| P35564 | Calnexin | 6.68 | 6.77 | 8.96 | 6.60 | 67.2 |
| P97461 | 40S ribosomal protein S5 | 12.68 | 16.18 | 15.78 | 16.18 | 22.9 |

**A.**

| **Accession** | **Description** | **Score Ferrous**  **Hb** | **Coverage Ferrous**  **Hb** | **MW**  **[kDa]** |
| --- | --- | --- | --- | --- |
| P08228 | Superoxide dismutase [Cu-Zn] | 6.89 | 38.96 | 15.9 |
| P24369 | Peptidyl-prolyl cis-trans isomerase B | 16.37 | 35.65 | 23.7 |
| P30681 | High mobility group protein B2 | 16.68 | 27.14 | 24.1 |
| Q923G2 | DNA-directed RNA polymerases I, II, and III subunit RPABC3 | 7.01 | 28.67 | 17.1 |
| P30416 | Peptidyl-prolyl cis-trans isomerase FKBP4 | 9.08 | 10.26 | 51.5 |
| P57784 | U2 small nuclear ribonucleoprotein A' | 8.14 | 15.29 | 28.3 |
| P14211 | Calreticulin | 2.56 | 4.57 | 48.0 |
| Q7TPR4 | Alpha-actinin-1 | 11.60 | 4.71 | 103.0 |
| P06745 | Glucose-6-phosphate isomerase | 17.87 | 13.62 | 62.7 |
| P27048 | Small nuclear ribonucleoprotein-associated protein B | 4.53 | 6.49 | 23.6 |
| P11103 | Poly [ADP-ribose] polymerase 1 | 7.92 | 4.54 | 113.0 |
| Q64727 | Vinculin | 30.40 | 10.13 | 116.6 |
| Q60668-4 | Isoform 4 of Heterogeneous nuclear ribonucleoprotein D0 | 7.39 | 6.27 | 30.8 |

**B.**

**C.**

| **Accession** | **Description** | **Score Ferric**  **Hb** | **Coverage**  **Ferric**  **Hb** | **MW**  **[kDa]** |
| --- | --- | --- | --- | --- |
| P97760 | DNA-directed RNA polymerase II subunit RPB3 | 8.24 | 19.64 | 31.3 |
| P97429 | Annexin A4 | 17.28 | 17.24 | 35.9 |
| Q9DB20 | ATP synthase subunit O, mitochondrial | 11.87 | 16.90 | 23.3 |
| P97496-2 | Isoform 2 of SWI/SNF complex subunit SMARCC1 | 15.54 | 6.70 | 120.0 |
| P97384 | Annexin A11 | 14.93 | 8.55 | 54.0 |

**Supplemental Table 1**. Autoantigens binding to (A) both ferrous Hb and ferric Hb, (B) ferrous Hb and (C) ferric Hb.
