## Supplemental Figures for "Hemoglobin drives inflammation and initiates antigen spread and nephritis in lupus"

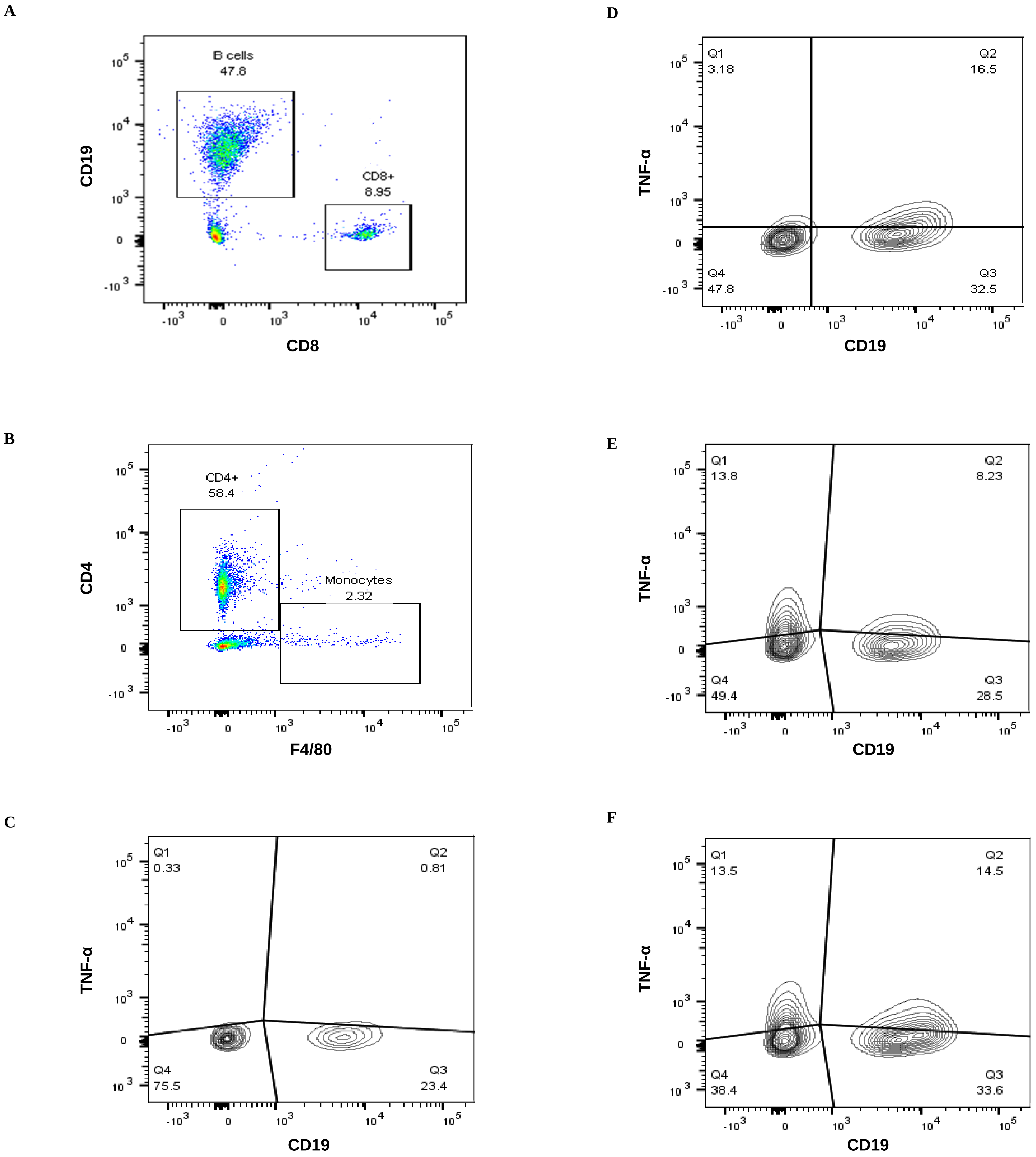


**Supplemental Figure 1. Flow cytometric analysis of intra-cellular cytokines.** (A) Representative gating strategies for B cells and CD8 T cells. (B) Representative gating strategies for CD4 T cells and monocytes. (C-F) Contour plots depicting intracellular TNF-α in B cells upon incubation of splenocytes derived from old NZM mice with (C) medium, or with medium supplemented with (D) LPS, (E) murine ferrous Hb, or (F) murine ferric Hb. Numbers in quadrants indicate percentages.

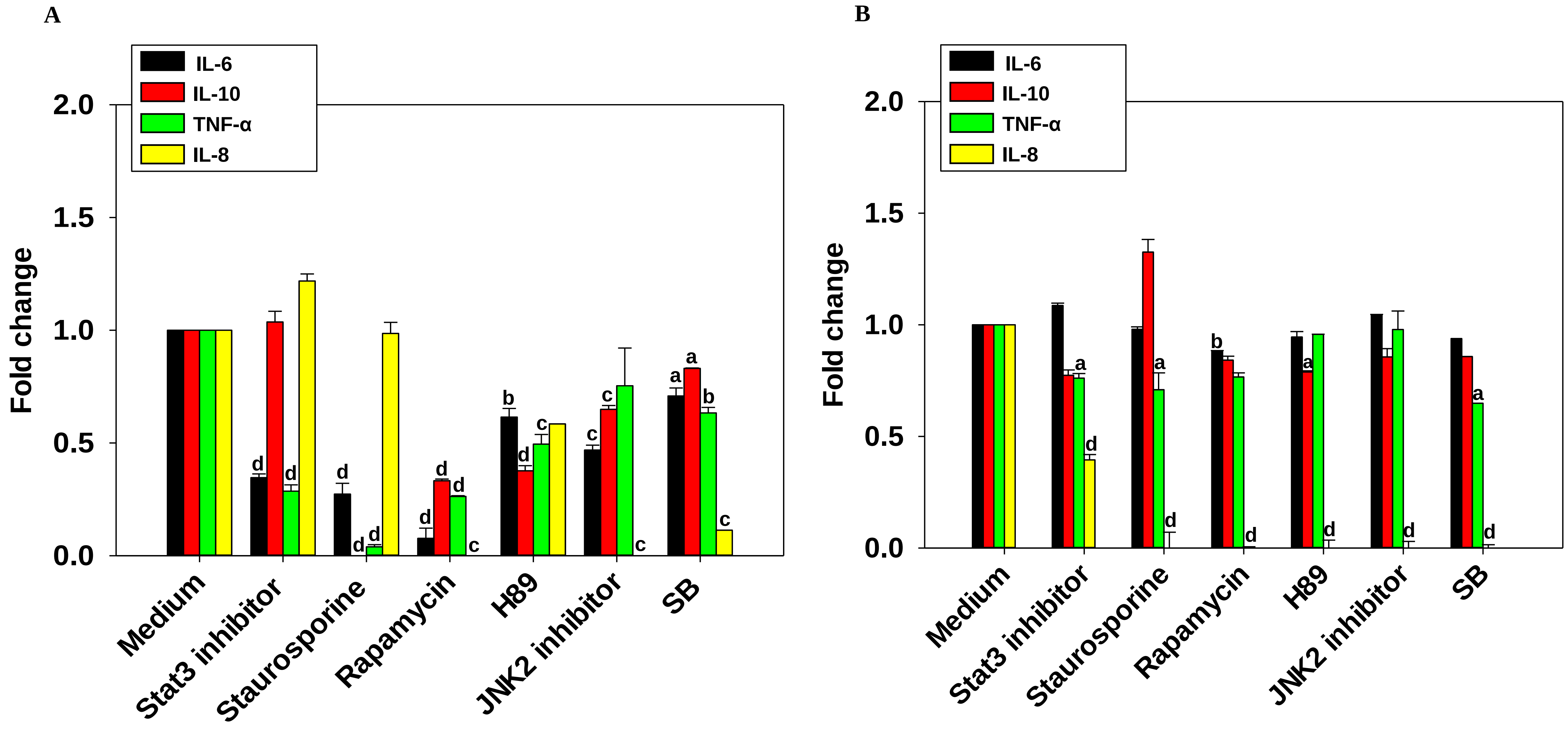


**Supplemental Figure 2. The effect of signaling inhibitors on Hb-induced cytokine secretion from splenocytes and pDCs.** (A) Splenocytes or (B) pDCs, isolated from old NZM mice (n=4) were incubated with different signaling inhibitors (or with an equivalent concentration of the vehicle control, "Medium") and then with medium supplemented with 0.5 µM murine ferric Hb. Levels of IL-6, IL-10, TNF-α and IL-8 in supernatants are depicted. Each experiment was carried out three times. Data is presented as "Fold Change" over Medium, and represents Mean ± SEM. $^{a}p<0.05$, $^{b}p<0.01$, $^{c}p<0.001$, $^{d}p<0.0001$ vs Medium.

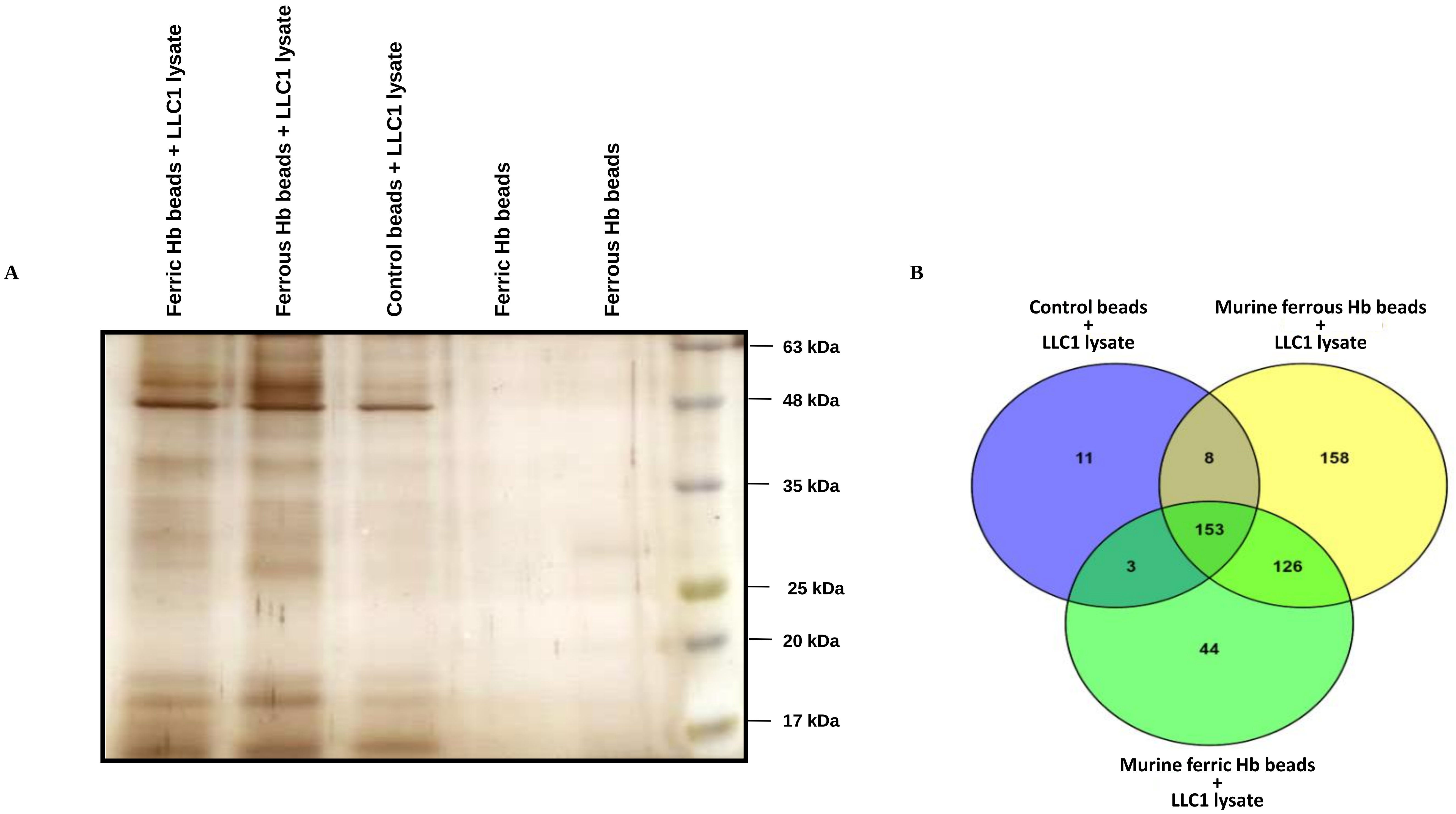


**Supplemental Figure 3. Assessment of Hb-self-moiety interaction.** (A) SDS-PAGE depicting self-moieties pulled down from LLC1 cell lysate by murine ferric Hb beads, murine ferrous Hb beads and control beads; murine ferric Hb beads and murine ferrous Hb beads, in the absence of LLC1 lysate, constituted additional controls. (B) Venn diagram indicating the number of self-moieties pulled down from LLC1 cell lysate (as determined by ESI-LC-MS/MS mass spectrometric analysis) by control beads, murine ferrous Hb beads and murine ferric Hb beads.

A

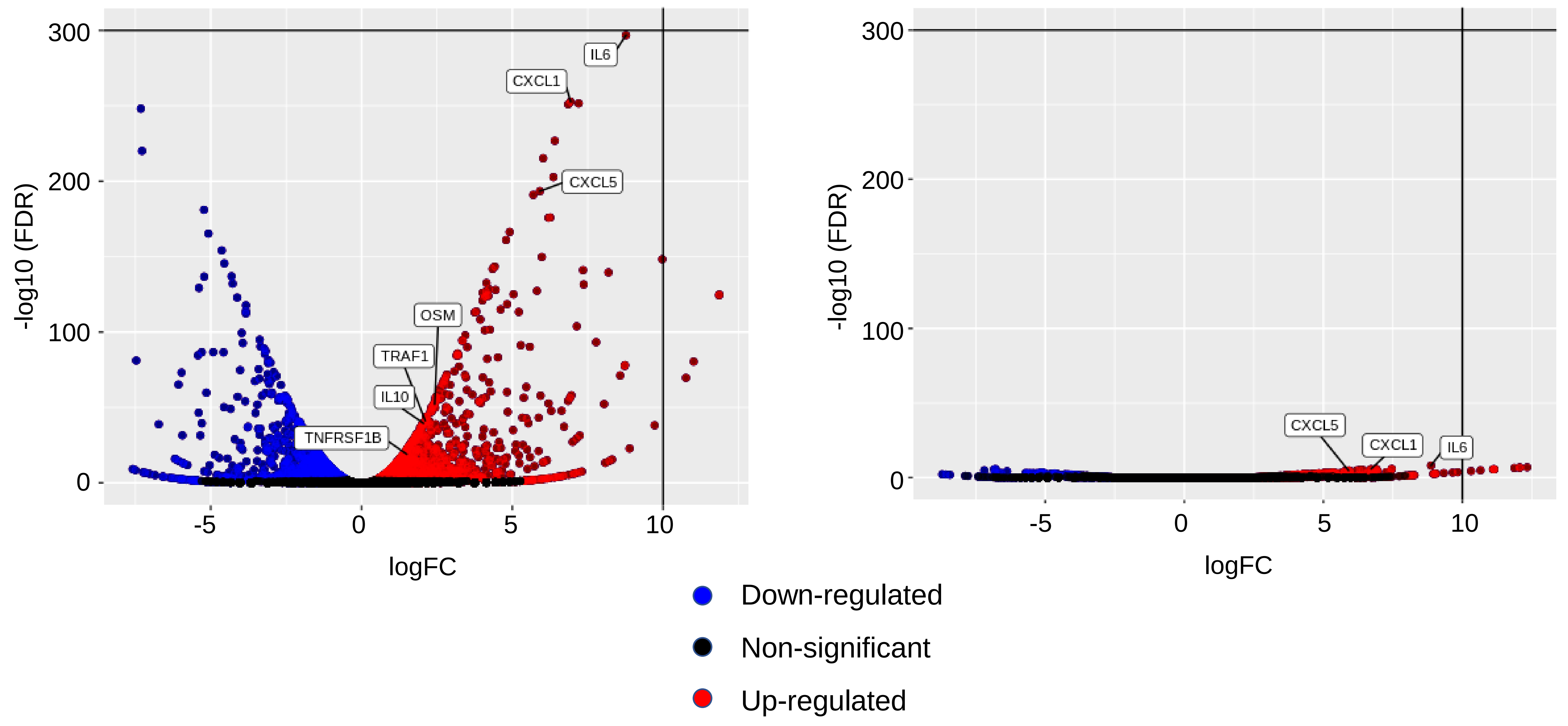


B

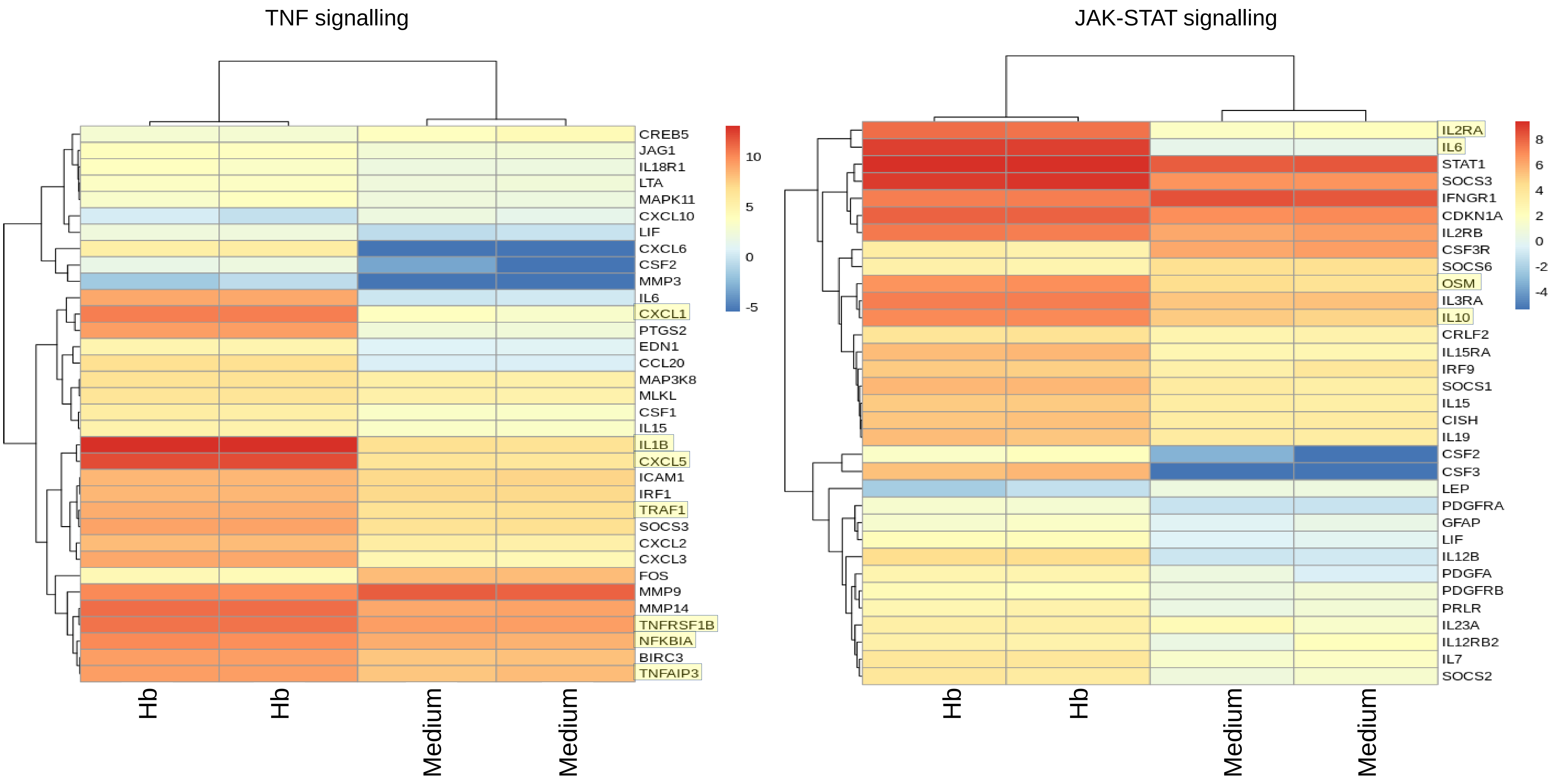


**Supplemental Figure 4. RNA-seq analysis on PBMC.** (A) Left panel: Volcano plot subsequent to the incubation of PBMCs (isolated from an SLE patient) with medium or with medium supplemented with 0.5 µM human Hb. Some mRNAs of potential interest have been highlighted. Right Panel: Volcano plot subsequent to the incubation of PBMCs (isolated from a healthy donor) with medium or with medium supplemented with 0.5 µM human Hb. Some mRNAs of potential interest have been highlighted. (B) Heat maps of mRNAs associated with the TNF signaling pathway (left panel) and the JAK-STAT signaling pathway (right panel) subsequent to the incubation of PBMCs (isolated from an SLE patient) with medium or with medium supplemented with 0.5 µM human Hb. Some mRNAs potential interest have been highlighted in yellow.
